# *In Silico* Benchmarking of Metagenomic Tools for Coding Sequence Detection Reveals the Limits of Sensitivity and Precision

**DOI:** 10.1101/295352

**Authors:** Jonathan Louis Golob, Samuel Schwartz Minot

## Abstract

High-throughput sequencing can establish the functional capacity of a microbial community by cataloging the protein-coding sequences (CDS) present in the metagenome of the community. The relative performance of different computational methods for identifying CDS from whole-genome shotgun sequencing (WGS) is not fully established.

Here we present an automated benchmarking workflow, using synthetic shotgun sequencing reads for which we know the true CDS content of the underlying communities, to determine the relative performance (sensitivity, positive predictive value or PPV, and computational efficiency) of different metagenome analysis tools for extracting the CDS content of a microbial community.

Assembly-based methods are limited by coverage depth, with poor sensitivity for CDS at < 5X depth of sequencing, but have excellent PPV. Mapping-based techniques are more sensitive at low coverage depths, but can struggle with PPV. We additionally describe an expectation maximization based iterative algorithmic approach which we show to successfully improve the PPV of a mapping based technique while retaining improved sensitivity and computational efficiency.

## Introduction

High throughput (or “next-generation”) sequencing has uncovered the hidden complexity of microbial communities living within and upon the human body, as well as the link between the human microbiome and health [1–4]. The taxonomic composition of a microbial community can be inferred by sequencing PCR amplicons spanning variable regions of a taxonomically informative gene (i.e. the 16S rRNA gene or the CPN60 gene)[5–8]. Alternatively, DNA recovered from a sample can be put through Whole-Genome Sequencing (WGS), which samples the complete genomic content of a sample via random fragmentation and sequencing [9]. WGS differs from amplicon sequencing by (1) providing genomic data from all organisms in a sample—not limited to any single domain of life; (2) enabling a high degree of taxonomic resolution which identifies the subspecies and strains present in a sample; and (3) generating a “functional” metagenomic profile of the protein coding sequences (CDS) that are present in a sample in addition to the organisms which contain those genes [10]. While the term “functional” can often be used to describe predicted metabolic pathways, in this case are limiting our scope to the identification of CDS without presupposing knowledge of any annotations.

There are three broad computational approaches used to generate an estimate of the functional metagenome (CDS content) of a microbial community from WGS reads: (1) The inferred taxonomic composition can be used to construct a custom database of protein-coding genes from the set of reference organisms detected in the sample (e.g. HUMAnN2, MIDAS) [11,12]. (2) *De novo* assembly, in which the WGS reads are combined into contigs, which can be further used to identify open reading frames (e.g. metaSPAdes, IDBA-UD) [13,14]. (3) The WGS reads can be directly mapped (aligned) to a closed reference of protein coding sequences (which is also a downstream component of HUMAnN2 and MIDAS).

Proteins can evolve by duplication events, truncation, homologous recombination, and other means that result in the sharing of highly conserved domains between otherwise distinct CDS [15]. As a result, mapping of reads to a closed reference of CDS is challenged by the fact that some reads may align equally well to multiple references: “multi-mapping” reads.

Metagenomic tools have been benchmarked extensively for their ability to determine the taxonomic composition of a microbial community [16–19]. The relative ability of metagenomic analysis approaches and tools to accurately infer the CDS catalog of a microbial community has yet to be established. Additionally, benchmarking efforts are often limited in their long-term utility by the practical challenges of repeating the computational analysis with the addition of newly available tools. We address this core challenge of benchmarking by implementing our analysis within a workflow management tool, Nextflow [20], which achieves a high degree of reproducibility by executing each component task within Docker containers, a portable and fixed computational environment.

Here we establish sensitivity and positive predictive value (PPV) of computational tools for determining the CDS content of a microbial community metagenome, using synthetic communities and reads generated *in silico* for which we know the true CDS content of the community. We establish that assembly-based approaches achieve a near-perfect PPV, but struggle with sensitivity for CDS at a low sequencing coverage depth. Mapping-based approaches are more sensitive, particularly at low coverage depths, but struggle with PPV. We introduce an expectation-maximization based approach for mapping based metagenomics that retains the sensitivity and improves the PPV of CDS calls close to that of assembly-based approaches.

## Materials & Methods

### Evaluating Computational Tools

All of the analytical steps for analyzing computational tools for CDS detection from metagenomes were executed within a single analytical workflow (‘evaluate-gene-detection.nf’) which can be downloaded from https://github.com/FredHutch/evaluate-gene-level-metagenomics-tools and executed via Nextflow. That analytical workflow follows this approach:

1. Simulate metagenomes (n=100)
  a. Randomly select host-associated genomes from NCBI/RefSeq (n=20). (A list of genomes from host-associated organisms is available in the supplemental materials.)
  b. Make a file with all of the CDS records from those genomes
  c. Assign sequencing depths for each genome from a log-normal distribution (mean=5x, std=1 log), with a maximum possible depth of 100x
  d. Make a file with the depth of sequencing for each CDS from step (1b) above
  e. Simulate reads from whole genome sequences via ART (paired-end read length 250bp, mean fragment length 1kb +/- 300bp)
  f. Interleave paired end FASTQ data
2. Run tools
  a. For assembly-based tools, perform assembly from paired end FASTQ data and predict CDS records from the resulting contigs
  b. For mapping-based tools, run the tool and then extract the FASTA for all detected CDS records
3. Perform evaluation
  a. For each tool, align the FASTA with all detected CDS records against the set of truly present CDS records (from step 1b)
    i. Prior to alignment, both sets of FASTAs are clustered at 90% amino acid identity to account for sets of homologous genes in the simulated metagenome
  b. Filter to the top hit for each detected CDS
  c. Assign each detected CDS as:
    i. True positive: The detected CDS is the mutual best hit for a truly present CDS
    ii. False positive: The detected CDS does not align against any truly present CDS
    iii. Duplicate: The detected CDS aligns against a truly present CDS, but is not the best hit (i.e. there are multiple non-overlapping detected CDS records that each align against a single truly present CDS).
  d. Calculate accuracy metrics:
    i. Sensitivity is calculated as the number of true positives (TP) divided by the number of true positives and false negatives (FN): TP / (TP + FN)
    ii. Positive Predictive Value is calculated as the number of true positives (TP) divided by the number of true positives and false positives (FP): TP / (TP + FP)
    iii. Uniqueness is calculated as the number of true positives (TP) divided by the number of true positives and duplicates (DUP): TP / (TP + DUP)

### FAMLI Implementation

FAMLI is available as an open source software package on GitHub at https://github.com/FredHutch/FAMLI. In addition, Docker images are provided at https://quay.io/repository/fhcrc-microbiome/famli to facilitate easy usage by the research community with a high degree of computational reproducibility. FAMLI can be run with the single executable “famli”, which encompasses:

1. Downloading reference data and query FASTQ files (supporting local paths, FTP, and Amazon Web Service (AWS) object storage)
2. Aligning query FASTQ files in amino acid space with DIAMOND
3. Parsing the translated alignments
4. Running the FAMLI algorithm to filter unlikely reference peptides and assign multi-mapping query reads to a single unique reference.
5. Summarizing the results in a single output file
6. Copying the output file to a remote directory (supporting local paths, FTP, and AWS object storage)

The help flag (“-h” or “—help”) can be used to print a complete list of options, including the flags used to run the filtering process starting from step 4 above..

### FAMLI Overall Approach

1. Align all input nucleotide reads in against a reference database of peptides; UniRef 90 was used for this study [21].
2. Calculate the coverage depth (CD) across the length of each reference, representing the number of reads aligning to each amino acid position of the reference.
3. Filter out any reference sequences with highly uneven coverage:

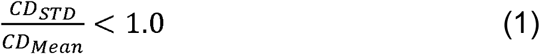 Where STD is standard deviation of per-base coverage values.
4. Calculate initial score for a given query coming from a subject using the alignment bitscores to weight the relative possibilities for a given query, normalizing the scores to total to 1 for a given query.
5. Iteratively, until no further references are pruned or a maximum number of iterations is reached:
  i. WEIGHTING and RENORMALIZING: The score of queries being from a subject from the prior iteration are weighted by the sum of scores for a given subject, and then renormalized to sum to 1 for each query.
  ii. PRUNING. Determine the maximum likelihood for each query. Prune away all other likelihoods for the query below a threshold.
6. Repeat filtering steps 2-3 using the set of deduplicated alignments resulting from step 4.

Here are some examples:

⍰ For reference A and reference B that both have some aligning query reads, if **there is *uneven* depth for reference A** but relatively even depth across reference B, then **reference A is removed from the candidate list** while reference B is kept as a candidate.
⍰ If query **read #1 aligns equally-well to reference A and reference C**, but **there is *2x more* query read depth for reference A as compared to reference C** across the entire sample, then **reference C’s alignment is removed from the list of candidates for query read #1**.

A more detailed description of the method is available in the supplemental materials. An interactive demonstration of our algorithm is available as a Jupyter notebook is available at https://github.com/FredHutch/FAMLI/blob/master/schematic/FAMLI-schematic-figure-GB.ipynb

### Comparison of FAMLI to HUMAnN2, SPAdes, Top Hit, and Top 20

The version of FAMLI presented in this paper is v1.3, which can be found at https://github.com/FredHutch/FAMLI/releases/tag/v1.3. FAMLI was executed in this analysis using a Docker image hosted at https://quay.io/repository/fhcrc-microbiome/famli with the tag v1.3 (sha256:25c34c73964f).

The version of DIAMOND used for translated nucleotide alignments in this analysis is DIAMOND v0.9.10 using a Docker image compiled from https://github.com/FredHutch/docker-diamond and available at https://quay.io/repository/fhcrc-microbiome/docker-diamond as v0.9.23--0 (sha256: 0f06003c4190).

Comparative analysis of the simulated communities used HUMAnN2 v0.11.1--py27_1, and all code used to run HUMAnN2 can be found in the GitHub repository https://github.com/FredHutch/docker-humann2 (v0.11.1--6), which is based on the BioBakery Docker image quay.io/biocontainers/humann2:0.11.1--py27_1. The Docker image used to run HUMAnN2 is available at https://quay.io/repository/fhcrc-microbiome/humann2 as v0.11.2--1 (sha256:d6426bda36ca).

The code used to run SPAdes is maintained by BioContainers and is available at https://quay.io/repository/biocontainers/spades as 3.13.0--0 (sha256:9f097c5d6d79). The code used to run megahit is maintained by BioContainers and is available at https://quay.io/repository/biocontainers/megahit as 1.1.3--py36_0 (sha256:8c9f17dd0fb1). The code used to run IDBA is maintained by BioContainers and is available at https://quay.io/repository/biocontainers/idba as 1.1.3--1 (sha256:51291ffeeecc). CDS were predicted from assembled contigs using Prokka as maintained by BioContainers (https://quay.io/repository/biocontainers/prokka) 1.12--pl526_0 (sha256:600512072486). The reference database used for the alignment-based analysis was UniRef90 (www.uniprot.org/uniref/) [16], downloaded on January 30^th^, 2018.

### Simulation of microbial communities

Synthetic microbial communities were simulated using ART (https://quay.io/repository/biocontainers/art) 2016.06.05--h869255c_2 (sha256:1cd93ed9f680) with paired-end reads, a read length of 250, mean fragment length of 1000, and fragment size standard deviation of 300. The abundance of each member of a given community was simulated from a log-normal distribution with a mean of 5x, standard deviation of 1-log, and maximum of 100x. Each community contains 20 distinct genomes.

## Results

### Sensitivity and specificity of metagenomics approaches

For each synthetic community, we cataloged the CDS present and compared these true positives to the reported CDS by each analytic method. For mapping-based methods, we allowed for duplicate calls (i.e. similar but distinct CDS sequences determined by the method to be roughly equally likely to be present). Comparing these CDS catalogs (true and inferred) we were able to calculate a positive predictive value (PPV; true positive / true positive + false positive), sensitivity (true positive / true positive + false negative), and uniqueness (true positive / true positive + duplicates). As shown in Figure 1, mapping-based approaches were more sensitive, particularly when the CDS has low coverage depth, at a cost of PPV and uniqueness.

**Figure 1:**
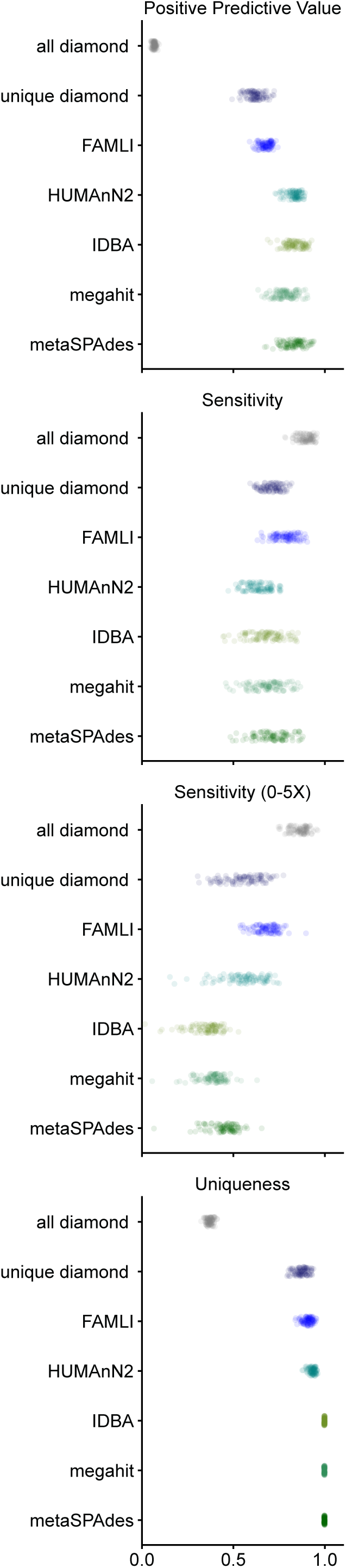
Positive predictive value (PPV), sensitivity, and uniqueness of CDS calls by metagenomic analysis approaches. The positive predictive value (true positive over true positive plus false positive), sensitivity (true positive over true positive plus false negative) both overall and subsetted to CDS with 0-5x coverage, and uniqueness (true positive over true positive plus duplicates) on a per-CDS basis with different analysis approaches.

The mapping all-hits approach is the simplest approach, accepting as present any CDS that had at least one aligning short-read sequence. While very sensitive, this approach had dismal PPV and uniqueness. A related mapping method is to restrict to CDS with at least one short read that maps uniquely to that CDS: Mapping - unique hits; this approach yielded balanced sensitivity and PPV. FAMLI uses an expectation maximization-based iterative approach (considering evenness of coverage and total coverage depth) and achieves somewhat superior sensitivity and PPV as compared to the Mapping - unique hit approach.

HUMAaN2 uses a hybrid approach, combining taxonomic identification, mapping of reads to reference genomes, and then using a mapping - all-hits like approach for the remainder of short reads that do not map to a genome. Our experimental set-up biases in favor of organisms with reference genomes. In this favorable set of circumstances, HUMAaN2 performs well with regards to PPV (superior to any of the tested mapping based approaches), sensitivity (similar at all depths and low-coverage depths, slightly inferior to mapping approaches) and with uniqueness.

Assembly based approaches have the advantage of near perfect uniqueness (with the assembly process itself resulting in convergence on a single CDS), and the best PPV. Sensitivity was inferior to mapping-based approaches, and varied by the coverage depth for a given CDS (Figure 2).

**Figure 2:**
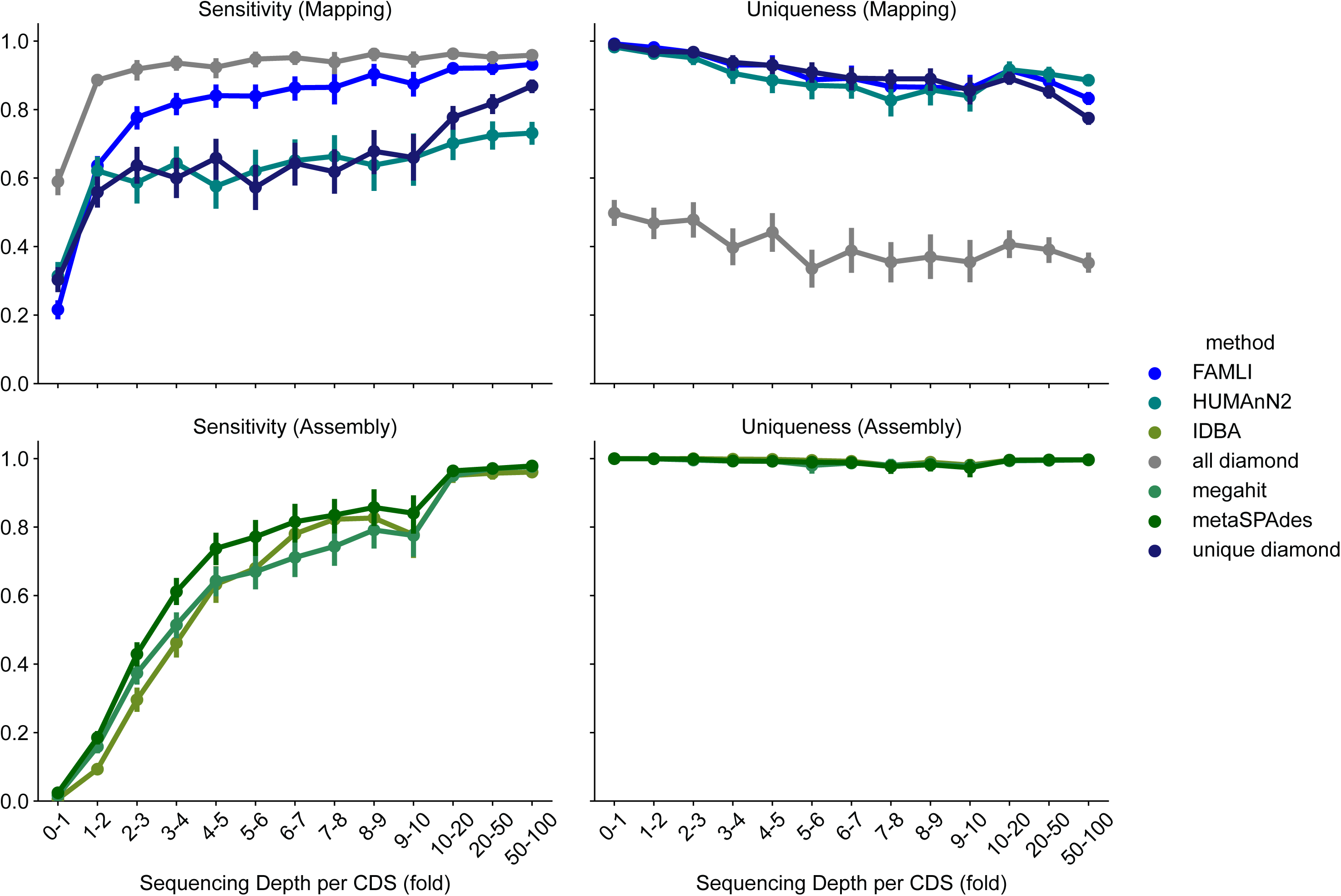
Sensitivity and uniqueness of CDS calls with respect to CDS coverage depth. Mapping based approaches are both more sensitive, and achieve a plateau of sensitivity at a lower coverage depth as compared to assembly-based methods.

### Short reads align equally well to multiple CDS

To better understand why mapping approaches, particularly mapping with acceptance of all hits, has poor sensitivity, we explored the role multi-mapping reads may be playing. To do so, three random unique CDS were selected and 120 simulated reads were generated for each CDS, resulting in a total of 360 simulated reads. These simulated reads were aligned against the UniRef100 database. Each read has only one true origin CDS.

To account for sequencing errors and poor representation in the reference database, we accepted alignments within a certain percentage of the best alignment for a given read. When we accept all CDS with an alignment within 10% identity (‘top-10’) of the best alignment for a read, 100,468 CDS are recruited for the 360 reads, an average of 279 (median of 268, minimum of 77 and maximum of 537) CDS recruited from UniRef100 per read (figure XXX D, Start).

When taking a more restricted approach, only recruiting CDS with an alignment to a read equivalent to the best hit, a total of 57,983 CDS are recruited, an average of 161 (median of 165, minimum of 1 and maximum of 384) equally well aligning reference CDS for each simulated read.

### The FAMLI approach can successfully cull multi-mapped reads

To establish the extent of the multimapping read problem, three random CDS were selected from UniRef100. One hundred and twenty simulated reads were generated from each CDS, and combined into one set of 360 paired reads; each of these reads had one true origin coding sequence.

We then used Diamond to align these 360 reads against UniRef100. Even after limiting to only alignments within equal in quality to the best hit, there were an average of 161 (median 165, min 1, and max 384) reference sequences tied with the best hit per read pair; when limited to alignments within 10% of the best identity, there was a mean of 279, median 268, minimum 77, and max 537 aligning subjects (references) per read pair.

To filter these alignments, we developed an iterative expectation maximization-based approach that considered both the evenness of coverage and total depth of coverage (weighted by alignment quality) of a candidate CDS in order to cull the vast excess of recruited CDS by the mapping approach, the FAMLI algorithm. Figure 3 shows the FAMLI algorithm applied to the top-10 alignments. Figure 3A shows the coverage (or read depth by base pair) for the three true positive CDS. After filtering for coverage evenness, Fig 3B shows the read-depth of some successfully filtered away references, as well as some references not present in the simulated sample that pass this evenness test. Figure 3C depicts the iterative pruning of alignments by likelihood, showing the candidate references for one query being successfully filtered down over ten iterations to a single reference CDS for the read (the true origin reference for this read). By the conclusion of the first evenness filtering, 908 references remain (for the true three); the 360 reads remain with an average of 271 (median of 267, minimum of 77, and maximum of 398) equally-well aligning reference CDS. By the conclusion of ten iterations, all reads are successfully assigned now to their true origin CDS (one reference CDS per read) (Figure 3D).

**Figure 3:**
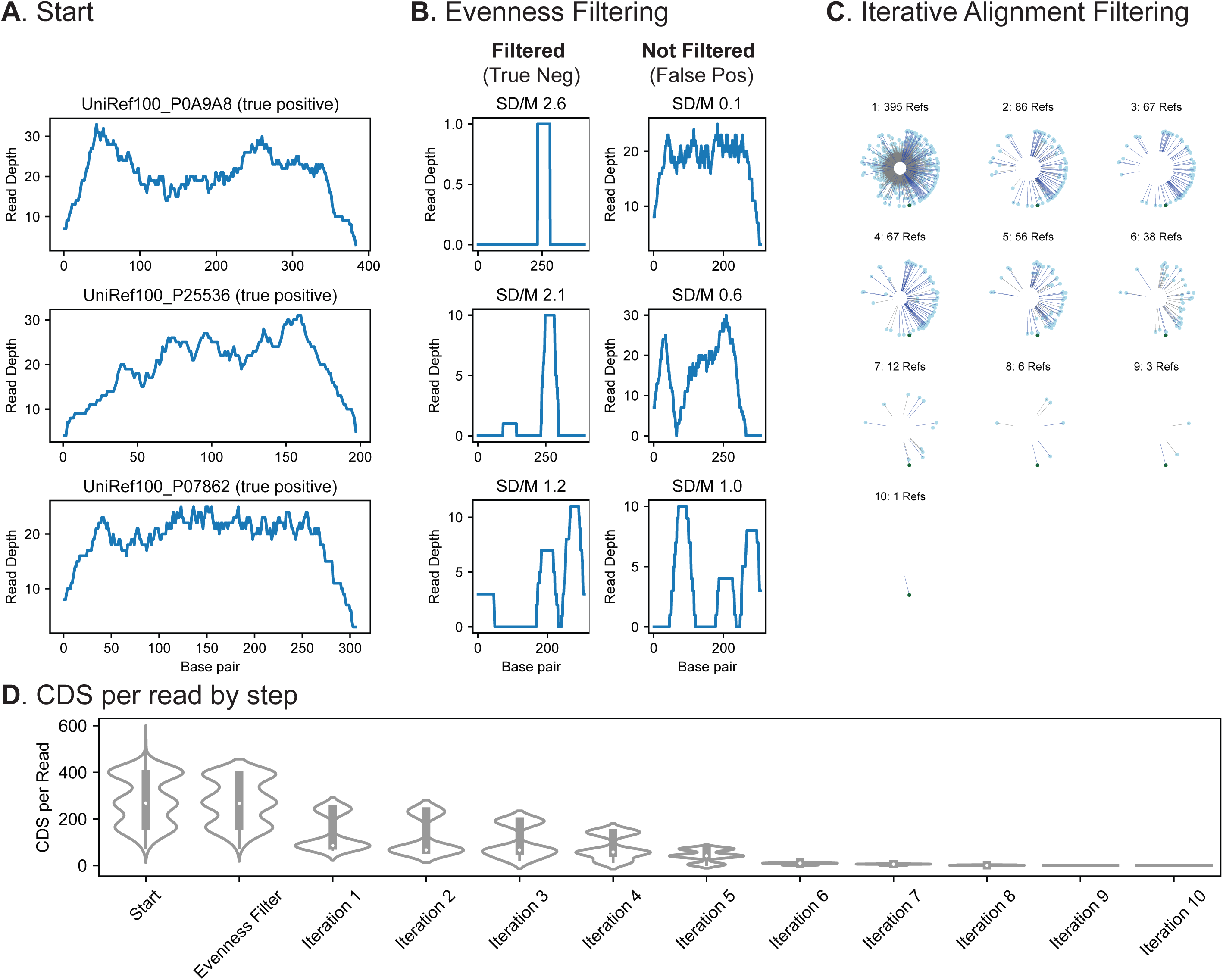
The problem of multiply-mapping short-reads, and the FAMLI algorithm schematized. Three hundred and sixty simulated reads were generated from three CDS. These simulated read was aligned against the UniRef100 database, and all CDS with an alignment within 10% identity of the best match were retained. A) The read-depth coverage of the three true peptides (top) B) Evenness filtering is used to remove the least likely to be present references from being considered. The left column is three randomly selected references that are successfully filtered at this step, the right three false references that are not filtered. C) The iterative likelihood-based filtering of one randomly selected synthetic read. Each circle represents one remaining aligned reference CDS for this read; the true positive origin reference is in dark green. The length of each line is proportional to the calculated score at this iteration. D)The number of CDS per read as a violin plot. After the tenth iteration, only one reference CDS (the correct) remains for this read.

## Discussion

Randomly fragmented shotgun sequencing of the metagenome of a microbial community offers the promise of inferring the functional capacity of the community by establishing the protein coding sequencing (CDS) present. CDS or gene-level metagenomics offers a more reproducible and mechanistic means of associating the state of the microbiome with functional outcomes in a host or environment [22]. Realizing this promise is predicated on having a reliable set of analytic tools for determining the CDS catalog of a microbial community.

Here we introduce and employ an approach for benchmarking the performance of different metagenome analysis tools for determining the CDS content of the metagenome. This benchmarking approach is implemented within a reproducible Nextflow workflow, and therefore should be relatively straightforward for other researchers to reproduce and augment as additional tools for CDS detection become available.

We found that assembly-based tools are limited by sensitivity, particularly at low read coverage. The association between the sensitivity to detect a CDS and the read coverage depth of the CDS is worrisome; the ability of these tools to detect a protein coding sequence is dependent upon community factors, including the relative abundance of the hosting organism, more so than other approaches.

Mapping-based approaches must address the problem of short-reads from metagenomes aligning equally well to large numbers of distinct CDS sequences. As evident in our simulated communities, the ratio of true to false positive alignments can be in the hundreds to one, resulting in dismal precision unless the alignments are culled or filtered. We suspect some of the limitations experienced by software attempting to use short reads to identify the functional genes encoded by microbial communities, described by [23], may be due to this multi-mapping read problem.

Here we demonstrate the magnitude of the problem of multiple-mapping of short reads to peptides, revealing a large number of equally-scored alignments; if one simply includes all peptides for which there is at least one short read that aligns equally as well as to any other peptide, the false positives outnumber true positives by an average of about 160:1.

We describe an algorithmic approach to correctly assign these multiply aligned WGS reads to the proper reference sequence, implemented as the open source software package FAMLI (Functional Analysis of Metagenomes by Likelihood Inference). With FAMLI, we are able to improve our precision (number of true positives divided by the sum of false and true positives) to about 80%; this performance is consistent over a range of community types. FAMLI is more efficient than *de novo* assembly at identifying protein-coding sequences present in a community with regards to both read depth and computational time. While FAMLI can be used as a standalone tool to identify protein-coding genes, it could also easily be used to enhance the precision of existing bioinformatics tools (e.g. HUMAnN2).

The hybrid approach of establishing which taxa are present and first mapping to reference genomes (e.g. HUMAnN2, MIDAS) has merit, and performed well from a sensitivity and positive predictive value perspective in our benchmarking approach. We note that our approach limits our synthetic communities to being those with reference genomes. This biases in favor of this hybrid approach. In the context of microbial communities with a high degree of novelty, we suspect performance would be poorer.

Thinking about the relative merits of reference-based (e.g. UniRef90) or reference-free (e.g. *de novo* assembly) analysis methods, one of our primary considerations was the efficiency of comparing results across large numbers of samples. While reference-free approaches are free by definition from the bias inherent in reference databases, that lack of common reference makes it extremely challenging to compare results between samples. For example, comparing a set of genes between N samples is an O(N^2) problem that scales exponentially with the number of samples. In contrast, by identifying proteins from a reference database (UniRef90), all results are inherently comparable without any additional computation (e.g. sequence alignment), in other words the complexity is O(1). Put simply, with *de novo* assembly (SPAdes) it is *much* more difficult to compare the results for 1,000 samples in contrast to just 10 samples, while for FAMLI or HUMAnN2 it is about the same.

## Supporting information

FAMLI approach details

## Acknowledgements

We would like to acknowledge funding support the Microbiome Research Initiative at the Fred Hutch, lead by David N Fredricks, for supporting this work and Dan Tenenbaum from the scientific computing group at the Fred Hutch for help with establishing computational resources for this manuscript.

