## Supplementary material for "*In Silico* Benchmarking of Metagenomic Tools for Coding Sequence Detection Reveals the Limits of Sensitivity and Precision": FAMLI approach details

### Filtering subject reference peptides by coverage evenness

Coverage evenness is calculated for every reference peptide sequence (Step 2, above). On a per-amino-acid basis, alignment depth is calculated using an integer vector. It is expected that the 5' and 3' ends of the reference will have trail offs, thus the depth vector is trimmed on both the 5' and 3' ends. A mean coverage depth and the standard deviation of the mean are calculated. The standard deviation is divided by the mean. Both based on the Poisson distribution and empirical efforts on our part, we set a threshold of 1.0 for this ratio as a cutoff of unevenness; **references with a coverage STD / MEAN ratio > 1.0 are filtered (Step 3, above)**.

### Defining alignment score

Let us consider the **score that a given query *q* is truly from a given reference r** considering all of the evidence from all of the queries in a sample. For the terms of this discussion, we will describe this as the **score** ($S_{qr}$) for a given assignment.

For this application we use the **bitscore**—an integrated consideration of the alignment length, number of mismatches, gaps, and overhangs—as a way of *initialization* for our score calculations: $Bitscore_{qr}$ is the quality of the alignment of query read *q* to reference peptide *r*.

We can use the bitscore of an alignment divided by the sum of bitscores for all the alignments for a given query sequence as a starting estimated **score** of any individual query *q* is truly derived from a reference *r*: $\mathbf{S_{qr}}$.

$$S_{qr} = \frac{Bitscore_{qr}}{\sum_{r=1}^{n_r} Bitscore_{qr}} \qquad (2)$$

Where $n_r$ is the number of references.

We then begin an iterative process. In each iteration:

1) We calculate the **total weight** for every reference r, $\mathbf{TOT_r}$ using the **score from the prior iteration,** $\mathbf{SP_{qr}}$.

$$TOT_r = \sum_{r=1}^{n_r} SP_{qr} \quad (3)$$

2) We calculate the new **score** that any individual query *q* is truly derived from a reference *r*, $\mathbf{S_{qr}}$, by first calculating a weighted score $\mathbf{W_{qr}}$

$$W_{qr} = SP_{qr} * TOT_r \quad (4)$$

and determine the new scores by renormalizing the scores for each query to 1.0.

$$S_q = \sum_{q=1}^{n_q} W_{qr} \quad (5)$$

$$S_{qr} = \frac{W_{qr}}{S_q} \quad (6)$$

3) The **maximum score for query q,** $\mathbf{Smax_q}$ is determined using the newly calculated scores.

$$Smax_q = max(S_{qr}\, for\ all\ r) \quad (7)$$

4) If the $S_{qr}$ falls below the scaled maximum score for query *q*, the **alignment is removed from consideration**:

For all query *q*, if

$$S_{qr} < scale * Smax_q \quad (8)$$

then $S_{qr}$ is set to zero.

By default the scale here is set to 0.9 (or 90% of the maximum score for query *q*).

**Iterations are repeated until no more alignments are culled** or a maximum number of iterations are reached.
